# A Movement-Independent Signature of Urgency During Human Perceptual Decision Making

**DOI:** 10.1101/2025.05.28.656559

**Authors:** Harvey McCone, Ciara A. Devine, Emmet McNickle, Jessica Dully, Anna C. Geuzebroek, David P. McGovern, Simon P. Kelly, Redmond G. O’Connell

## Abstract

How does the brain adjust its decision processes to ensure timely decision completion? Computational modelling and electrophysiological investigations have pointed to dynamic ‘urgency’ processes that serve to progressively reduce the quantity of evidence required to reach choice commitment as time elapses. To date, such urgency dynamics have been observed exclusively in neural signals that accumulate evidence for a specific motor plan. Across three complementary experiments, we show that a classic ERP component, the Contingent Negative Variation (CNV), also traces dynamic urgency but exhibits unique properties not observed in effector-selective signals. Firstly, it provides a representation of urgency alone, growing only as a function of time and not evidence strength. Secondly, when choice reports must be withheld until a response cue, the CNV peaks and decays long before response execution, mirroring the early termination dynamics of a motor-independent evidence accumulation signal. These properties suggest that the brain may use urgency signals not only to expedite motor planning but also to hasten cognitive deliberation. These data demonstrate that urgency processes operate in a variety of perceptual choice scenarios and that they can be monitored in a model-independent manner via non-invasive brain signals.

**Significance Statement:** Computational models suggest that, when decisions are time-constrained the brain progressively lowers the amount of evidence it requires to reach choice commitment, thus increasingly sacrificing accuracy for timely decision completion. To date, neurophysiological investigations have identified signatures of these ‘urgency’ effects exclusively in areas of the brain that plan the decision-reporting actions. Here, we characterise a human electroencephalogram signature of urgency that exhibits several novel properties: it traces the urgency component of the decision and terminates upon choice commitment even when the decision-reporting action is deferred until later. These observations suggest that urgency can serve to hasten the deliberation process and not just the movements that a decision entails.

## Introduction

As the motorist fast approaching a fork in the road can appreciate, making a quick decision can be just as important as making an accurate one. In recent years, research has made significant progress toward identifying the key algorithmic adjustments that allow observers to strike an appropriate balance between speed and accuracy during decision making. The dominant computational accounts hold that perceptual choices are formed through a process of accumulating sensory evidence into a decision variable that triggers commitment upon reaching a bound ^1–3^. This bound can be strategically lowered in order to shorten deliberation times when speed must be prioritized or raised to allow more evidence to accrue when accuracy is paramount ^4,5^. In standard models, these decision bound adjustments are made prior to deliberation and maintained until commitment is reached^6^. However, in other model variants, the bounds can dynamically ‘collapse’ over time, reducing the amount of evidence required to reach commitment as time elapses ^7–9^. Although many behavioural modelling studies have shown that such dynamic adjustments are often not necessary to include in a model to comprehensively fit behaviour ^10,11^, neurophysiological investigations have consistently indicated that both static and dynamic bound adjustments are implemented when choices must be made to a strict deadline. Under speed pressure, choice-selective motor planning signals exhibit elevated starting levels of activity, and, in addition to the evidence-driven component of build-up during stimulus evaluation, exhibit an evidence-independent, time-dependent ‘urgency’ build-up component ^12–18^. These urgency components effectively implement a bound collapse by progressively reducing the quantity of additional evidence required for motor planning signals to reach their action-triggering thresholds ^12–18^.

Thus far, such urgency components have been exclusively observed within neural populations that represent the evolving decision variable as a specific motor plan, such as monkey lateral intraparietal area (LIP) when choices are reported via saccades ^12,15^, and human limb-selective mu/beta (8-30 Hz) activity when choices are reported via manual button pushes ^16–18^. This apparent effector-dependence raises the question of whether urgency signals are encoded exclusively within motor planning circuits, or whether there are signals in the human brain that encode urgency on its own, and may play a more general, movement-independent role in expediting choice commitment. In the present study, our efforts to characterize a candidate signature of urgency in the human electroencephalogram (EEG) yielded new insights into this question.

Our analyses centred on a well-studied component of the human event-related potential known as the Contingent Negative Variation (CNV). The CNV is a slow negative-going potential observed over fronto-central scalp sites that is classically examined during the interval prior to the onset of an expected stimulus that informs an action contingency ^19–21^. Several well-established characteristics of the CNV identify it as a candidate marker of urgency. First, the CNV has been linked to various aspects of temporal processing, including stimulus and response anticipation ^22–25^. Second, the amplitude of the CNV predicts the speed with which individuals make motor responses in target detection and choice response tasks ^26–30^. Third, slow scalp potentials like the CNV have been linked to the regulation of cortical excitability thresholds ^31–33^. Finally, when making perceptual decisions, the magnitude of the CNV measured prior to stimulus onset has been shown to increase with experimenter-induced time pressure and correlates with trial-to-trial variations in model-estimated response caution (i.e. distance between the decision variable’s starting point and bound ^34^). While these observations suggest that the CNV may reflect processes that adjust decision boundaries prior to decision making, the extent to which the CNV might also trace bound adjustments that are implemented *during* deliberation has not yet been investigated.

In the present study, we sought to comprehensively examine the CNV’s evolution both before and during decision formation under varying regimes of speed versus accuracy emphasis. In so doing, we reveal that the CNV exhibits key regime-dependent features in its dynamics that are consistent with those expected of evidence-independent urgency and which mirror those observed in effector-selective decision signals and simulations of an urgency model fit to behaviour. In addition, we demonstrate that the CNV peaks and returns towards baseline long before delayed manual reports are executed consistent with a motor-independent representation of urgency.

## Results

### Experiment 1. The CNV builds at a time-dependent rate and is modulated by speed emphasis but not evidence strength

In Experiment 1, 25 subjects performed a two-alternative forced-choice contrast discrimination task in which they reported whether the left-or right-tilted lines in a compound overlay pattern had a greater contrast by pressing a button with the thumb of the corresponding hand (Fig 1a). The task was performed in six blocks of alternating Regimes in which subjects were instructed to emphasise either the speed or accuracy of their choices. At the outset of each trial, participants were presented with a cue instructing them to ‘Be accurate’ or ‘Go fast’. The stimulus then appeared with the gratings at equal contrast for an initial lead-in period of 800ms after which the choice evidence was presented i.e. one grating increased and the other decreased in contrast. This lead-in was used to ensure that visual potentials evoked by the stimulus onset would not obscure decision-related neural activity. As expected, choices were significantly faster (*F*(1,24) = 84.27, *p* < 0.001, η^2^_p_ = 0.78) and more accurate (*F*(1,24)=53.81, *p* < 0.001, η^2^_p_ = 0.69) on trials with strong relative to weak evidence (Fig 1, b and c). In addition, choices were significantly faster (F(1, 24) = 32.82, p < 0.001, η^2^_p_ = 0.58) and less accurate (*F*(1, 24) = 14.18, p < 0.001, η^2^_p_ =0.37) in the Speed regime (Fig 1, b and c). We also observed a significant Regime (Speed vs Accuracy) by Evidence Strength (Strong vs Weak) interaction for RT (*F*(1,24) = 10.69, *p* = 0.003, η^2^_p_ = 0.31) arising from a larger effect of Difficulty in the Accuracy regime. Plotting accuracy as a function of RT (“Conditional Accuracy Function”, Figure 1c) revealed that accuracy levels peaked at intermediate RTs in both Regimes and then declined for longer RTs, consistent with dynamic urgency progressively lowering the evidence accrued at commitment.

**Figure 1:**
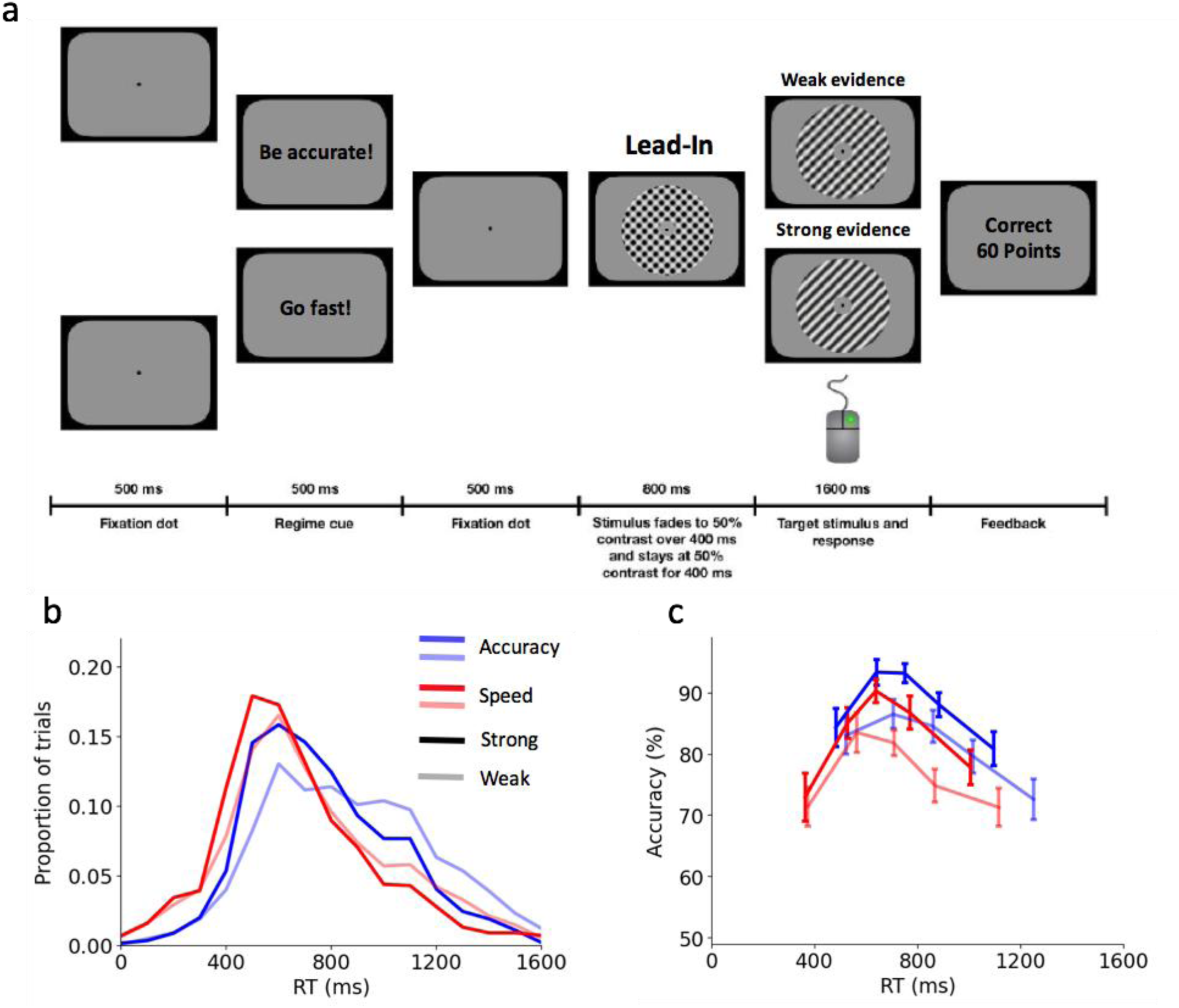
a) Task schematic. b) Response time distributions plotted for each Regime (Accuracy = blue, Speed = red) and Evidence Strength level (Strong = darker lines, Weak = lighter lines). c) Conditional accuracy functions depicting accuracy as a function of response time. Error bars represent standard error of the mean.

Previous work has established that mu/beta activity exhibits a progressive amplitude reduction during deliberation at a rate that scales with evidence strength and reaches a stereotyped amplitude contralateral to the chosen hand movement immediately prior to response execution ^35,36^. In addition, mu/beta exhibits adjustments to its starting levels and build-up rate in response to experimental manipulations of time pressure that accord with the operation of a dynamic urgency process ^16–18^. Here, in line with these previous studies, we found that mu/beta signals contralateral to the chosen response reached a stereotyped amplitude immediately prior to response execution (−150ms to -50ms from response) that did not vary across Regimes (Figure 2a; *F*(1,24) = 1.54, *p* = 0.23, η^2^ = 0.06) or RT bins (Figure 2b; F(3,72) = 0.78, *p* = 0.51, η^2^ = 0.03). Motor preparation for both alternatives began to increase in advance of evidence presentation in both Regimes, consistent with there being an evidence-independent, time-dependent component to its build-up. In addition, there was significantly greater mu/beta desynchronisation by the time of evidence onset (−100ms to +100ms) in the Speed regime (Main effect Regime: *F*(1,24) = 6.45, *p* = 0.02, η^2^ = 0.21; Figure 2a). Pre-response mu/beta desynchronisation over ipsilateral sites was also unaffected by Regime (*F*(1,24) = 0.05, *p* = 0.82, η^2^ = 0.002; Figure 2a and b) but increased significantly as a function of RT bin (*F*(3,72) = 4.12, *p* = 0.009, η^2^_p_ = 0.15; Figure 2a) consistent with the influence of a dynamic urgency process driving motor preparation signals for both alternatives towards their thresholds as time elapses ^17^. Together, these mu/beta results provide neurophysiological evidence of the operation of a dynamic urgency process.

**Figure 2.**
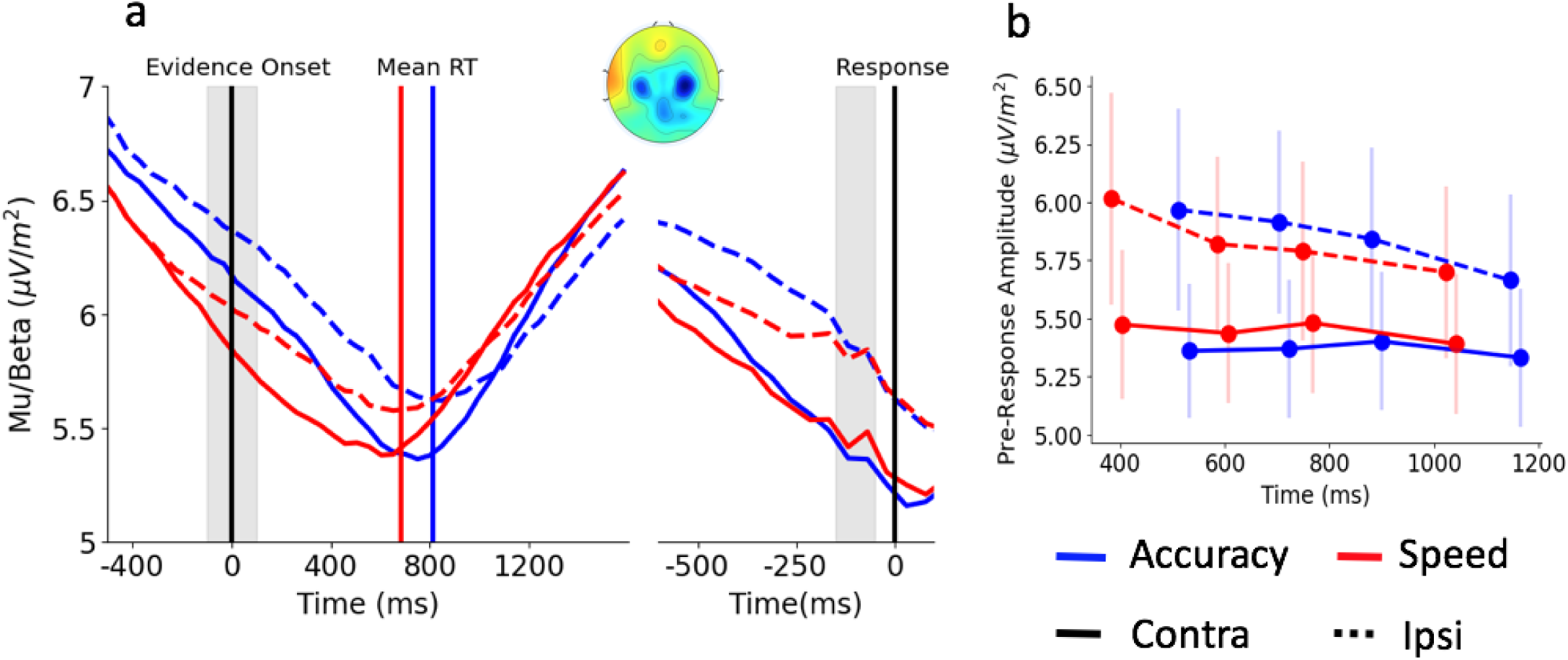
Mu/beta (10-30Hz) motor preparation signals contralateral (solid lines) and ipsilateral (dashed lines) to the response-hand plotted as a function of Regime a) Motor preparation signals plotted relative to evidence onset and response time. Motor preparation builds in advance of evidence onset, with a larger amount of build-up in the Speed Regime. Irrespective of Regime, the contralateral signals reach a fixed threshold at response. Shaded areas represent the time windows used for calculating amplitude at evidence onset and prior to response. Topography represents the difference in amplitude between evidence onset and response. b) Contralateral mu/beta amplitude at response (solid lines) did not vary as a function of Regime or RT Bin. Ipsilateral mu/beta amplitude at response (dashed lines) did not vary as a function of Regime, but decreased with RT, indicating increased preparation of the unchosen response.

To further verify the decision strategies used by participants, and to simulate urgency time courses for comparison with the CNV, we also fit a variant of the drift diffusion model^1^ to the behavioural data of the Accuracy and Speed Regimes (Figure 3a and b). This model conformed to the standard drift diffusion model with the exception that the accumulation process was initiated during the lead-in period (−500ms from evidence onset) and the bounds could be dynamically adjusted over time. Early accumulation was included since previous investigations using similar tasks and stimuli have identified that a substantial amount of sensory noise is accumulated when lead-in periods are introduced prior to evidence onset ^37^. In order to capture dynamic urgency effects, the change in the decision bound as a function of time was specified by a three parameter Weibull function^10^ that onset alongside evidence accumulation. This model identified decision bound adjustments that accorded with the mu/beta observations above: the bounds were lower at evidence onset in the Speed Regime, and collapsed in an approximately linear fashion over time in both Regimes (Table 1, Figure 4c).

**Figure 3:**
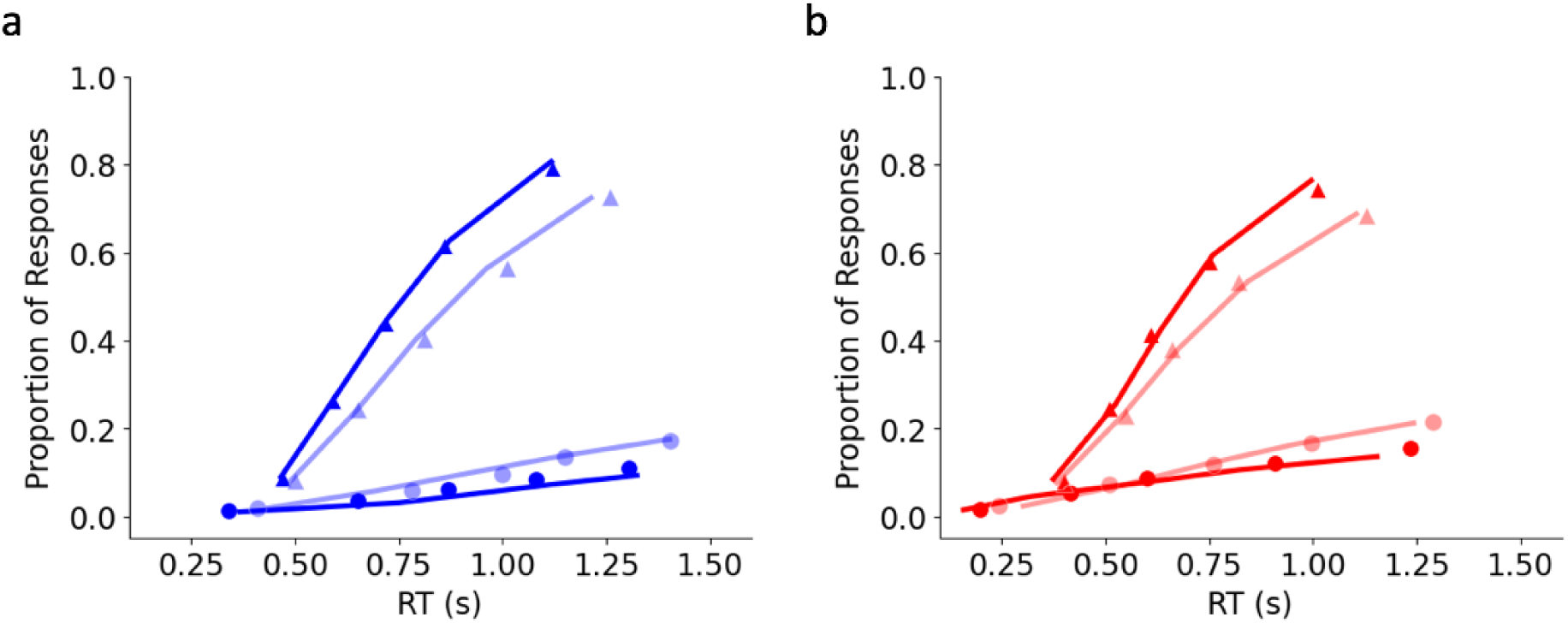
Model fits. Each panel shows the fit of the model (lines) to the observed cumulative proportion of correct (triangles) and error (circles) responses for strong (bold) and weak (faded) Evidence Strength, plotted as a function of RT. Fits to the Accuracy Regime are in blue (a), and fits to the Speed Regime are in red (b).

**Table 1:** Parameter values and G^2^ for model fitted to each Regime. The Shape and Time Parameters refer to aspects of a nonlinear function of time describing bound collapse.

|  | Bound | Drift Rate<br>Strong | Drift Rate<br>Weak | Ter | Shape<br>Parameter | Time<br>Parameter | Asymptotic<br>Boundary | G<sup>2</sup> |
| --- | --- | --- | --- | --- | --- | --- | --- | --- |
| Accuracy Regime | .41 | .32 | .21 | .28 | 1.18 | 3.90 | 1.48 | 36 |
| Speed Regime | .52 | .30 | .20 | .29 | .70 | 2.34 | 1.39 | 36 |

**Figure 4:**
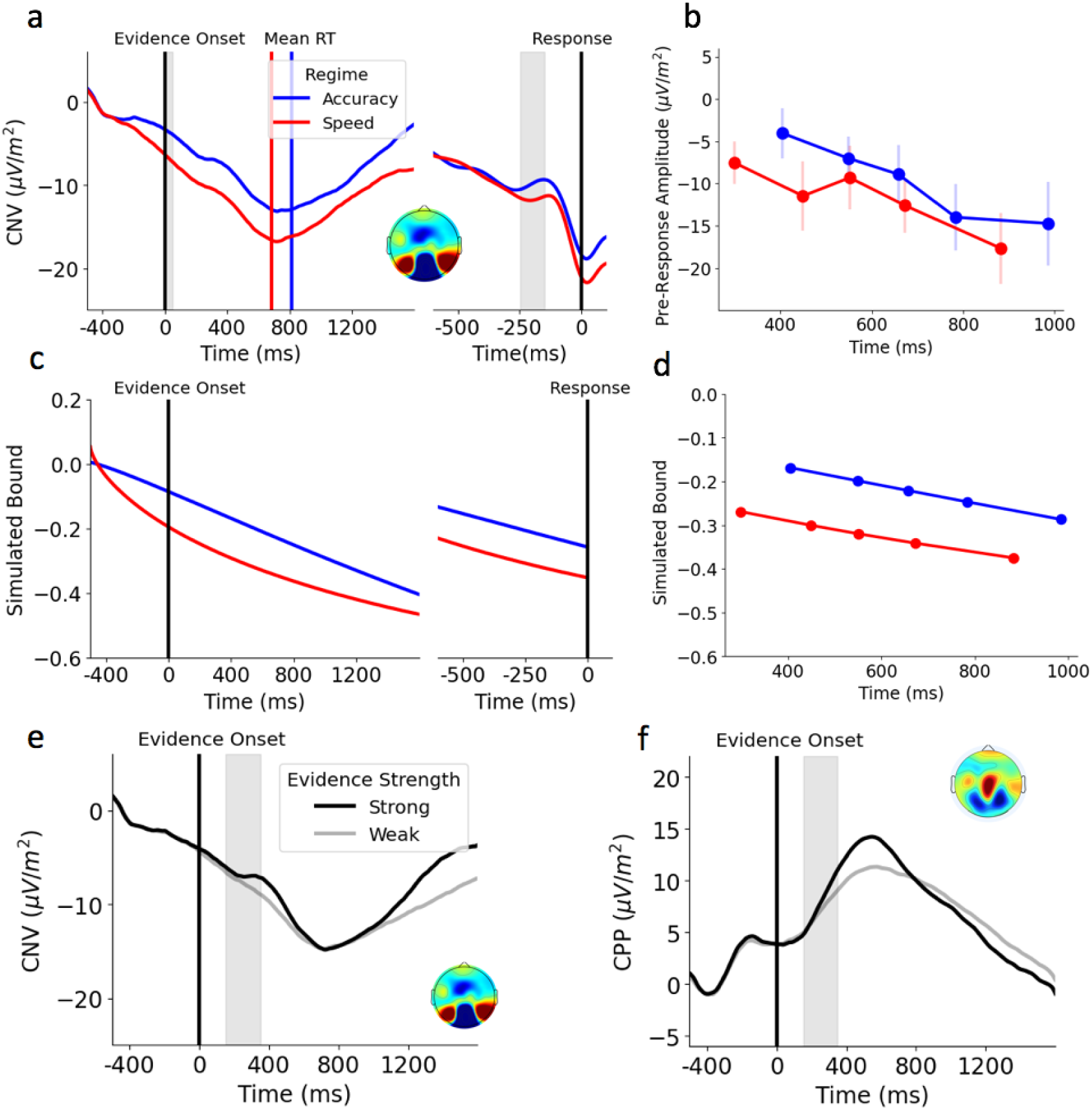
a) CNV plotted as a function of Regime relative to evidence onset and response. Shaded areas represent the measurement window for signal amplitude. Topography in this panel represents the grand-average slope at each electrode at evidence onset. b) CNV amplitude prior to response at response got larger (more negative) as a function of RT, but did not differ across Regimes. c) Change in decision bound over time simulated from Weibull Bound Collapse Model (left panel aligned to evidence onset, right panel aligned to response. A pre-evidence onset (−500ms to -400ms from evidence onset) baseline subtraction was applied to the simulated decision bound in order to facilitate comparison with the CNV. d) Simulated bound height plotted as a function of RT Bin. e) Evidence-onset aligned CNV plotted as a function of Evidence Strength. The slope of the CNV shortly after evidence onset (shaded portion) did not vary with Evidence Strength. f) Evidence-onset aligned CPP plotted as a function of Evidence Strength. As expected, the slope of the CPP was steeper for Strong sensory evidence. Topography in this panel represents grand-average amplitude prior to response (−150ms to -50ms).

Consistent with previous observations of the CNV, a fronto-central negative-going signal emerged during the lead-in period between stimulus and evidence onset (Figure 4a). The signal continued to build monotonically during evidence presentation, reaching its peak at the time of the choice report (Figure 4a). As the Regimes in all three experiments reported in this manuscript were administered in separate blocks of trials, our analyses are not well-suited to measuring static adjustments in the CNV due to the requirement for baseline subtraction of the single-trial data (note that Regime effects on mu/beta, which does not require baseline correction, are evident at the outset of the epoch in Figure 2). To provide some scope to observe anticipatory dynamics as evidence onset approached, we applied baseline correction using the interval -500ms to -400ms in advance of the evidence. In line with the mu/beta and model-predicted bound adjustments, the CNV had a significantly larger (more negative) amplitude at evidence onset in the Speed compared to the Accuracy regime (*F*(1,24) = 5.48, *p* = 0.03, η^2^ = 0.19; Figure 4a and b). Its pre-choice amplitude also increased as a function of RT in both regimes, (*F*(4,96) = 3.45, *p* = 0.01, η^2^ = 0.13; Figure 4b). The CNV reached larger pre-choice amplitudes in the Speed Regime although the difference did not reach statistical significance (*F*(1,24) = 1.98, *p* = 0.17, η^2^_p_ = 0.08; Figure 4b) and there was no significant Regime by RT Bin interaction (*F*(4,96) = 0.31, *p* = 0.87, η^2^ = 0.01; Figure 4b). These dynamics are highly consistent with those observed for mu/beta and the CNV reproduces the qualitative pattern of dynamic adjustment and RT-dependence seen in the model simulated data.

A defining feature of dynamic urgency components specified in extant decision models is that their build-up is independent of sensory evidence strength (Hanks et al., 2014; Hawkins et al., 2015). To test this, we compared the effect of Evidence Strength (Strong vs Weak) on the build-up rate of the CNV. For comparison, we conducted the same analysis on a well-established neural signature of evidence accumulation, the Centro-Parietal Positivity (CPP) for which evidence-dependent buildup is a central characteristic ^38^. For this analysis, we measured the build-up rates of both the CNV and CPP in an evidence-aligned, rather than response-aligned, window in order to ensure that any purely time-dependent build-up components would be matched across Evidence Strength levels. We measured signal slopes between 150ms to 350ms from evidence onset (shaded area in Figure 4e & f), removing trials with RTs occurring before the end of this time window in order to prevent any post-response dynamics from impacting on our measurements. For this analysis, we multiplied the CNV slopes by -1 to give them the same polarity as the CPP. We observed a significant Signal (CPP vs CNV) by Evidence Strength (Strong vs Weak) interaction (*F*(1,24) = 18.97, *p* < 0.001, η^2^_p_ = 0.44; Figure 4e and f) driven by the fact that the CPP’s build-up rate was reliably faster when evidence was strong (*F*(1,24) = 24.41, *p* < 0.001, η^2^_p_ = 0.50; Figure 4f), while the build-up of the CNV was not significantly affected by Evidence Strength (*F*(1,24) = 2.97, *p* = 0.10, η^2^_p_ = 0.11; Figure 4e). This functional dissociation also serves to rule out the possibility that the CNV is a dipole of the CPP.

### Experiment 2: The CNV is movement-independent

In order to further examine the CNV’s functional characteristics, we interrogated another dataset in which participants made dot motion direction (left vs right) discriminations under varying levels of difficulty (0%, 5%, 10% and 20% coherence) and time pressure, the latter of which was manipulated across three different regimes: A) 1200ms response deadline; B) 1800ms response deadline; C) Responses withheld until the appearance of a cue at 1800ms. The inclusion of the delayed response Regime enabled us to examine whether the time course of the CNV is more closely aligned with evidence accumulation (indexed by the CPP) or response execution. As in Experiment 1, the conditional accuracy functions (Fig 5c) exhibited a marked decline in accuracy for later RTs in each of the three Regimes. As expected, trials with stronger evidence were associated with significantly increased choice accuracy (*F*(3, 87) = 398.47, *p* < 0.001, η^2^_p_ = 0.93) and faster RTs (*F*(3, 87) = 99.92, *p* < 0.001, η^2^_p_ = 0.78). Additionally, there was a main effect of Regime on accuracy (*F*(2, 58) = 37.97, *p* < 0.001, η^2^_p_ = 0.57; Fig 5c) and RT (F(2, 58) = 2907.52, p < 0.001, η^2^_p_ = 0.99; Fig 5b and c). Accuracy and RT were highest in the Delayed Response condition and the lowest in the 1200ms deadline condition (Fig 5c). There was also a significant interaction between Regime and Evidence Strength for Accuracy (*F*(6, 174) = 4.16, *p* < 0.001, η^2^_p_ = 0.13), driven by the absence of any Regime effects on choice accuracy on trials with 0% coherence. There was also a significant interaction between Regime and Evidence Strength for RT (*F*(6, 174) = 65.47, *p* < 0.001, η^2^_p_ = 0.69), as the effect of Evidence Strength on RT was weaker in the Delayed Response Regime.

**Figure 5:**
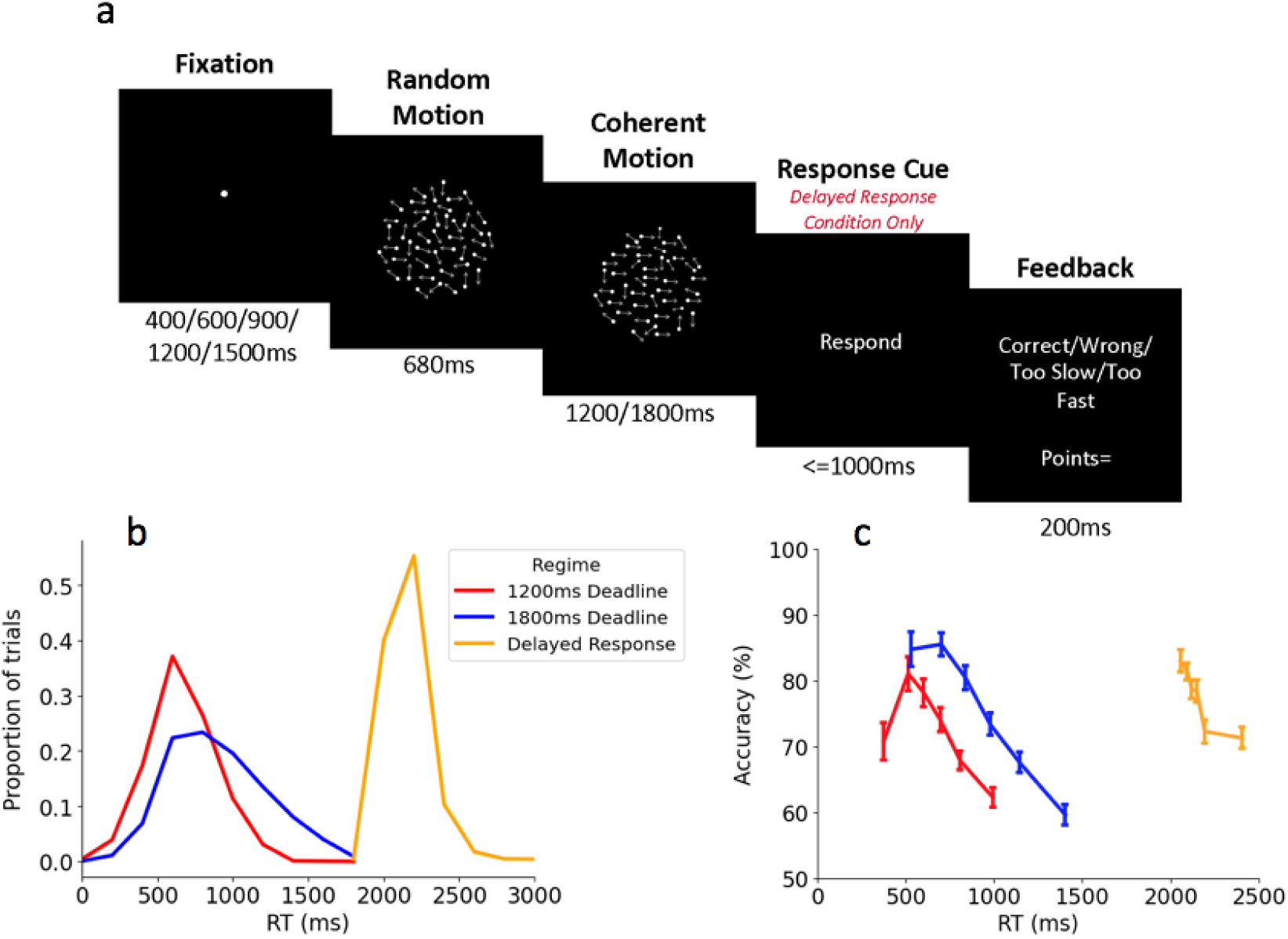
a) Schematic of RDM task. In the two Deadline Regimes, participants had to respond within 1200ms and 1800ms from evidence onset respectively. In the Delayed Response Regime, participants had to wait until the presentation of a response cue at 1800ms before making their response. b) RT histograms for each Regime. c) Accuracy plotted as a function of RT Bin for the three Regimes. RT in the Delayed Response Regime was measured relative to response cue onset. Error bars represent the SEM.

As in Experiment 1, motor preparation began to build in anticipation of evidence onset and there was a significant main effect of Regime on its amplitude at evidence onset (*F*(2, 58) = 20.88 , *p* < 0.001, η^2^_p_ = 0.42). This effect was primarily driven by the Delayed Response Regime, which had significantly less mu/beta desynchronisation than the other two Regimes (Fig 6a), while there was no significant difference in the level of mu/beta desynchronisation between the two Deadline Regimes (Fig 6a; M_diff_ = 0.05, 95% CI [-0.29, 0.38 ], *t* = 0.35, *p*_bonf_ = 1.00 , *d* = 0.02). In addition, contralateral mu/beta reached a fixed threshold at response (−150ms to -50ms from response) across the two Deadline Regimes (*F*(1, 29) = 0.17, *p* = 0.69, η^2^_p_ = 0.01), and RT Bins (*F*(3, 87) = 2.04 , *p* = 0.11, η^2^_p_ = 0.07), while ipsilateral mu/beta desynchronisation did not vary across Regimes (F(1, 29) = 0.19 , p = 0.66, η^2^_p_ = 0.01) but increased with RT Bin (*F*(3, 87) = 5.22, *p* = 0.002 , η^2^_p_ = 0.15).

**Figure 6:**
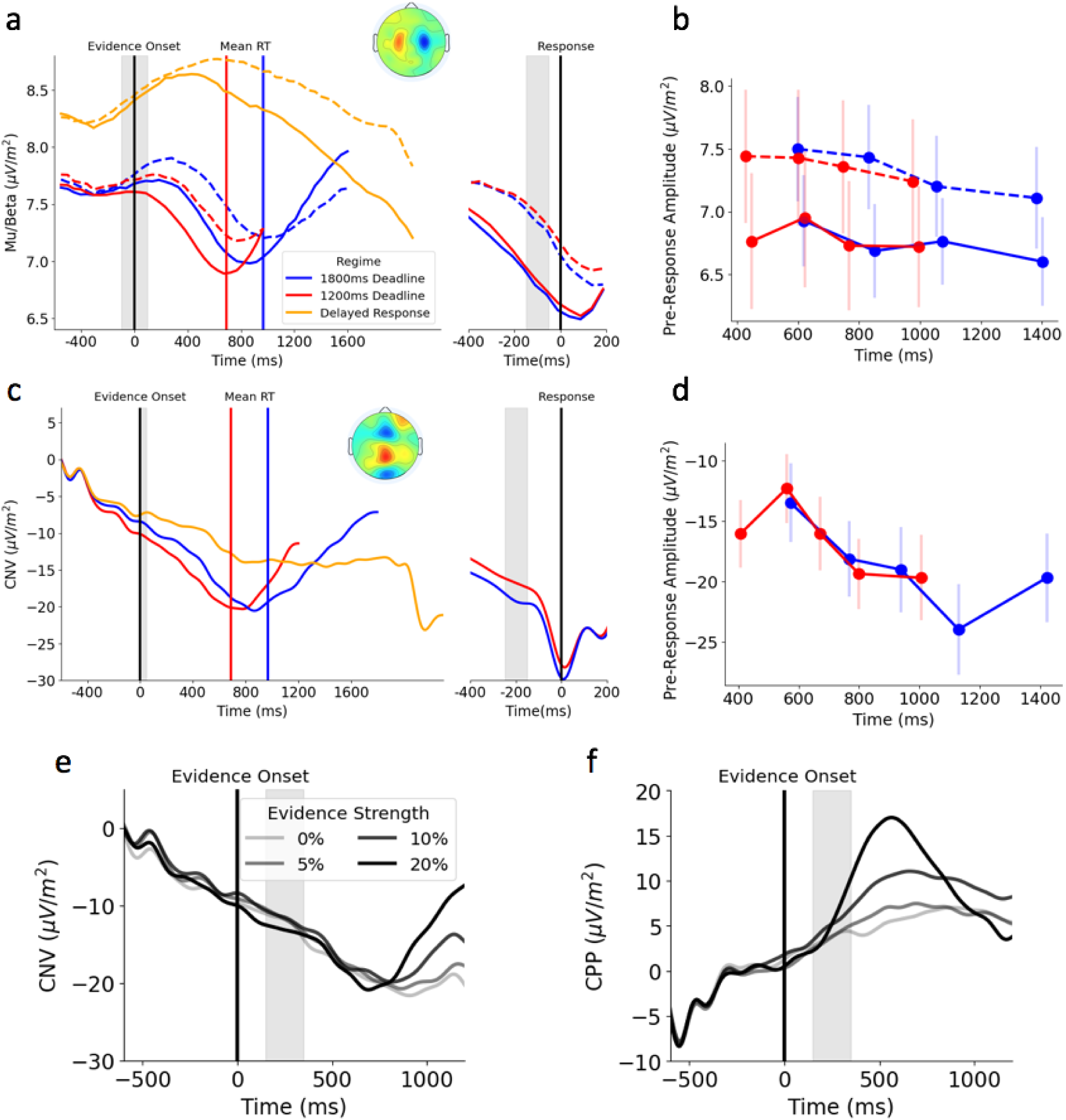
a) Mu/beta plotted as a function of Regime relative to evidence onset and response. Shaded areas represent the measurement window for signal amplitude. Topography in this panel represents grand average pre-response amplitude after subtracting activity for left responses from right responses b) Group mean (black lines and large markers) and participant mean (grey lines and small markers) mu/beta amplitude prior to response. c) CNV plotted as a function of Regime relative to evidence onset and response. Shaded areas represent the measurement window for signal amplitude. Topography in this panel represents the grand average pre-response amplitude. d) Group mean (black lines and large markers) and participant mean (grey lines and small markers) CNV amplitude prior to response. e) Evidence-onset aligned CNV and f) CPP plotted as a function of Evidence Strength.

As in Experiment 1, the CNV began to build in anticipation of evidence onset and exhibited larger amplitudes at evidence onset when greater time pressure was applied (Fig 6c) although the effect of Regime did not reach statistical significance (*F*(2, 58) = 2.87, *p* = 0.06, η^2^_p_ = 0.09). As in Experiment 1, its amplitude at response got larger (more negative) as a function of RT Bin (*F*(4, 116) = 6.25, *p* < 0.001, η^2^_p_ = 0.18) but did not vary across Regimes (*F*(1, 29) = 1.78, *p* = 0.18, η^2^_p_ = 0.06; Fig 6d). We repeated the analysis from Experiment 1, to test whether CNV build-up was evidence-independent, using only trials from the two speeded response conditions where it was possible to identify and remove trials with RTs falling within or before this measurement window. As in Experiment 1, we found a significant Signal by Evidence Strength interaction (*F*(3,87) = 3.74, *p* = 0.014, η^2^_p_ = 0.11) driven by a significant effect of Evidence Strength on CPP (*F*(3,87) = 10.36, *p* = 0.0007, η^2^_p_ = 0.26) that was absent for the CNV (*F*(3,87) = 0.09, *p* = 0.95, η^2^_p_ = 0.003; Fig 6e and f).

We examined the time course of the CNV in the Delayed Response condition to determine whether it peaked alongside choice commitment or response execution. For this analysis, we exclusively examined trials with 20% coherence on the basis that participants were likely to have made their decision well before the appearance of the response cue in this condition. We found that the CNV peaked well in advance of response execution (Fig 7a), returning toward baseline shortly after the termination of evidence accumulation indexed by the CPP peak (Fig 7b). In contrast, motor preparation signals exhibit a steady build-up that peaks at response execution (Fig 7c). This indicates that the CNV resembles an urgency signal driving the decision process to choice commitment, rather than an anticipatory signal that is preparing for an upcoming response cue.

**Figure 7:**
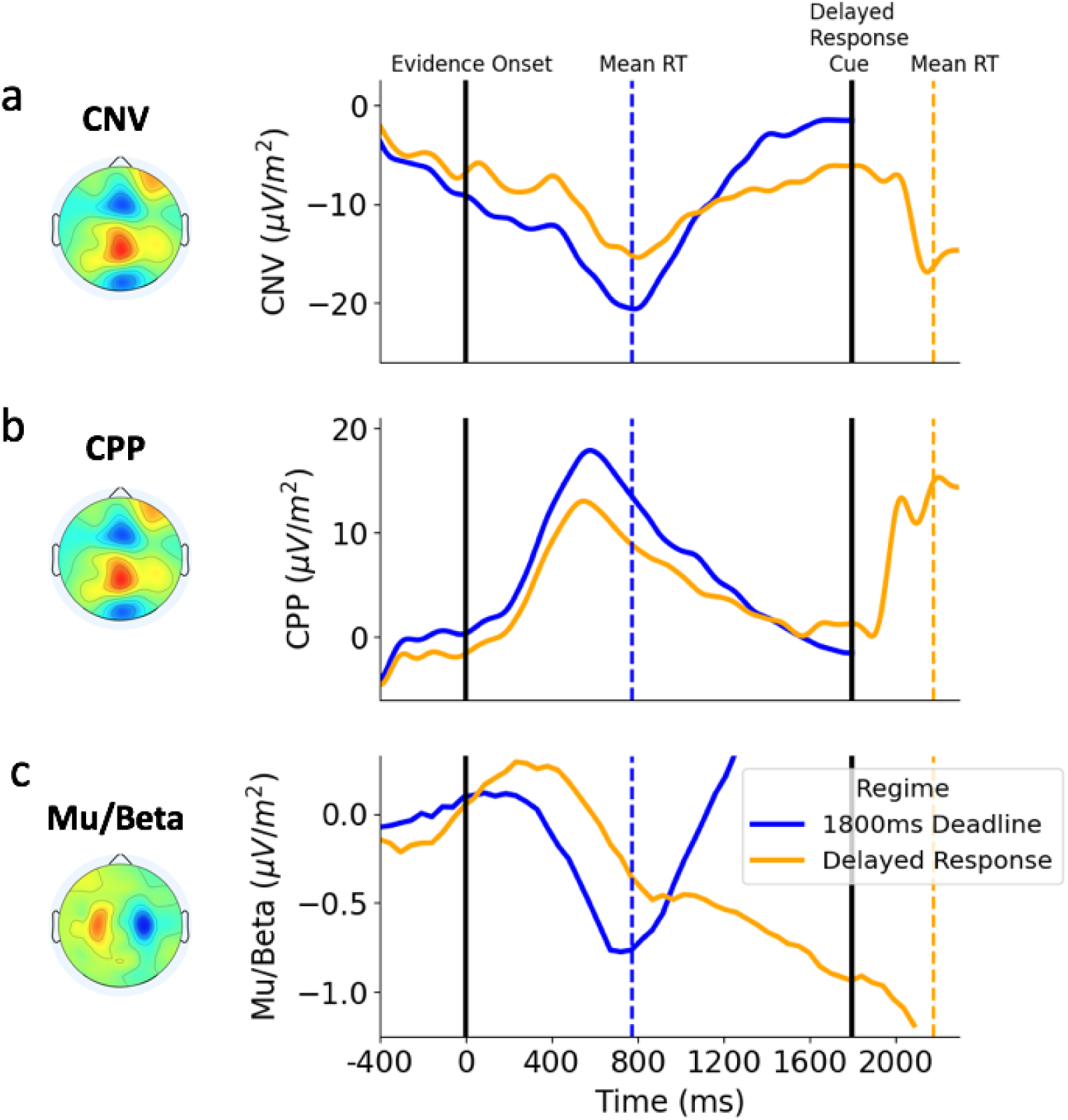
CNV (a), CPP (b) and Mu/Beta (c) in the 1800ms (blue) and Delayed Response (orange) Regimes. Each trace is plotted exclusively from trials with 20% coherence level. Rather than building until response execution, the CNV peaks and resolves to baseline alongside the termination of evidence accumulation, indexed by the CPP peak. To allow better comparison to the ERPs, Mu/Beta in this plot is baseline corrected by subtracting its amplitude at the start of the epoch from all time points.

### Experiment 3: CNV traces dynamic urgency adjustments in continuous monitoring settings

Our recent work has examined the decision policies that observers implement when faced with continuous monitoring scenarios in which targets can appear at unpredictable times ^39^. In this context, we found that mu/beta-indexed motor preparation increases steadily in anticipation of the onset of a target during the inter-target interval, indicative of increasing urgency as targets became more probable ^39^. Given that both signals traced urgency in a similar fashion in Experiments 1 and 2, we wanted to assess whether the time course of the CNV matched mu/beta motor preparation over the relatively long inter-target periods in continuous monitoring scenarios (up to 8 seconds).

Twenty-three participants (14 females, mean age: 22.5 years, age range: 18-29 years) performed a perceptual choice task (Fig 8a) in which they monitored a continuously presented random dot motion stimulus for targets defined by 2 second intervals of leftward or rightward coherent motion that had unpredictable onsets (inter-target intervals varying pseudorandomly between 3.6 and 8.4 seconds). Participants reported the direction of detected motion with speeded left or right mouse button clicks. The task was performed under alternating Speed and Accuracy Regime blocks (see Methods for details). For the following analyses focusing on CNV and mu/beta time courses over the ITI, we averaged across Regimes as there was no difference in CNV amplitude or slope between Regimes (Regime-specific waveforms are provided in Figure S2 and a full reporting of Regime effects on behaviour and neural indices of decision making will be reported in a forthcoming paper). Consistent with increasing urgency as time elapses, longer ITIs were associated with a higher False Alarm rate (Fig 8d; *F*(2, 44) = 14.85, *p* < 0.001, η^2^_p_ = 0.40), faster Mean RT to the following target (Fig 8b; *F*(2, 44) = 37.81, *p* < 0.001, η^2^_p_ = 0.63), and lower rate of missing those targets (Fig 8c; *F*(2, 44) = 5.92, *p* = 0.01, η^2^_p_ = 0.21). Following the approach of Geuzebroek et al., (2023), we examined the time course of mu/beta (10-30Hz) activity reflecting motor preparation over the ITI. As in this previous study, motor preparation initially decreased prior to the earliest possible ITI before steadily increasing. As in Experiments 1 and 2, the CNV time course bore a striking correspondence to that of mu/beta, exhibiting the same inverted U pattern throughout the ITI (Fig 8e) and we found that the mu/beta time course was highly predictive of the CNV time course (R^2^ = 0.87, *F*(1, 302) = 1931, *p* < 0.001, *r* = 0.93).

**Figure 8.**
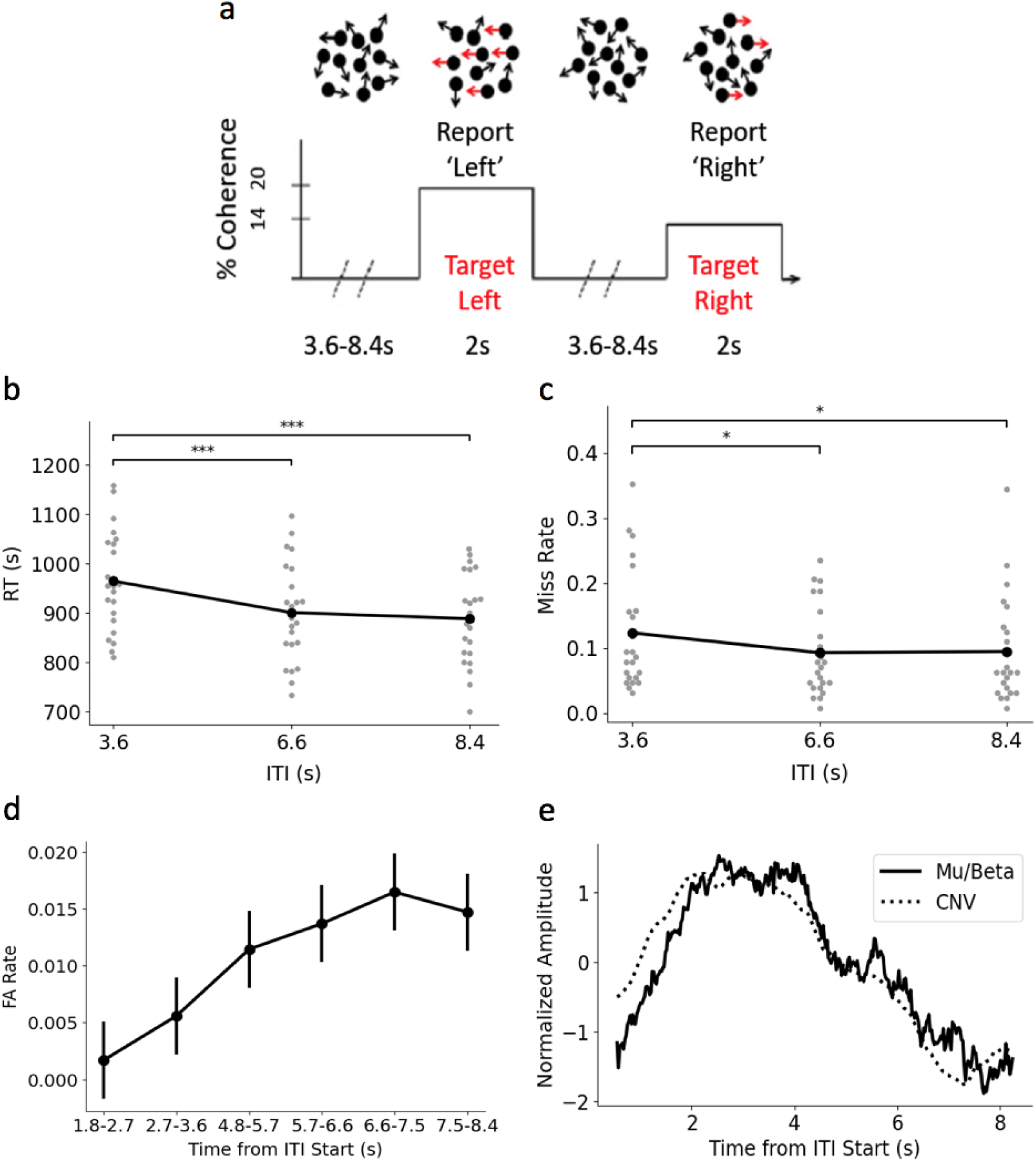
a) Continuous monitoring random dot motion task. Participants monitored a patch of randomly moving dots for intermittent periods of leftward or rightward motion with variable onsets (3.6s, 6.6s, 8.4s), reporting motion direction with left and right mouse clicks. In line with an increase in urgency, longer ITIs were associated with faster RTs (b) and decreased Miss rates (c). Asterisks indicate bonferroni corrected p-values for pairwise comparisons (* = p<.05, *** = p < .001). d) Additionally, False Alarm rate increased as the ITI progressed. False alarm rates were calculated within two consecutive 0.9s bins prior to the end of each of the three ITIs. e) In line with these behavioural trends, around the time of the earliest possible target onset (3.6s), mu/beta (solid line) gradually desynchronised (i.e. motor preparation increased) as the ITI progressed. The CNV (dotted line) followed a similar trajectory to mu/beta.

## Discussion

In the present study, we have shown that the CNV builds throughout decision formation at an evidence-independent rate, with a time course that aligns with decision bound adjustments identified through behavioural modelling and analysis of motor preparation signals. Moreover, we established that these effects are observed across multi-second timescales in continuous monitoring scenarios, as well as in the discrete-trial tasks that are most commonly studied in perceptual decision making research. In so doing, our results highlight several important functional dissociations between the CNV and motor preparation signals. First, whereas mu/beta motor preparation signals build as a function of both evidence strength and elapsed time, the CNV builds only as a function of elapsed time. Second, whereas contralateral motor preparation exhibited a threshold-crossing relationship with response execution in all conditions, the CNV’s peak is time-locked with the culmination of evidence accumulation and not response execution when these two events are separated in time. In addition, our results and interpretation of the CNV as a movement-independent urgency signal provide a potentially unifying account that can reconcile previous functional accounts that have implicated it in processes such as stimulus anticipation, response preparation, decision bound adjustment and time interval estimation.

A key focus of recent research on perceptual decision making has been to characterise the distinct algorithmic elements and neurophysiological processes that underpin decision formation ^38^. The apparent motor-independence of the CNV raises interesting questions regarding at what stage or stages in the sensorimotor hierarchy urgency is implemented. Previously, electrophysiological signatures of urgency had exclusively been found in areas associated with the preparation of specific effectors, taking the form of an evidence-independent, time-dependent gain in build-up rate ^12,15,17^. Thus, in the same way that the CPP represents a movement-independent representation of the same evidence accumulation process indexed by effector-selective decision signals ^40^, the CNV may represent a movement-independent representation of the urgency components that are also indexed by effector-selective signals. The data from the delayed response condition of Experiment 2 suggest that motor-independent urgency signals may play a role in regulating the termination of evidence accumulation processes when decisions do not entail immediate action. However, future work involving causal manipulations of CNV generators would be required to definitively test this proposal.

Electrophysiological signatures tracing distinct elements of the decision making process have emerged as key tools for evaluating and constraining computational models of perceptual decision making ^3,41^. Such signatures are especially useful in cases where competing models make similar predictions for behavioural data despite having different architectures. As such, in addition to previously identified signatures of urgency, the CNV could aid efforts to inform computational models and further characterise urgency adjustments across different contexts. For instance, signals like the CNV can be used to test for dynamic bound adjustments in datasets that may not have sufficient behavioural data for detailed modelling (e.g. clinical studies), or where such adjustments may be too subtle to detect through conventional modelling approaches (e.g. due to low trial numbers). However, it should be noted that while the CNV provides a means of tracing dynamic urgency adjustments, the baseline correction that must be applied to the signal can reduce its ability to capture static adjustments at the outset of a decision. Indeed, although we observed a significant difference in CNV amplitude at evidence onset between Regimes in Experiment 1, the effect did not reach statistical significance in Experiment 2 despite following a similar trend of a larger amplitude under increased time pressure. Task designs in which time pressure is manipulated and cued on a trial-to-trial basis would facilitate the detection of starting-level adjustments in the CNV.

The present results also have implications for long standing debates surrounding the functional role of the CNV. Recent work examining time estimation has converged on the consensus that the CNV represents a general anticipatory signal indexing the readiness to process or respond to upcoming task-relevant events (for reviews see ^23–25^). For example, the CNV exhibits highly similar adjustments during time interval reproductions to those we observed in our perceptual choice data with elevated starting levels and faster build-up rates in the lead-up to shorter reproductions ^42^. This suggests that the dynamics exhibited by the CNV in the present study are not particular to perceptual decisions. Instead, they may reflect a more general process that tracks the passage of time and the anticipation of upcoming task-relevant events, in a manner that can be scaled and calibrated to meet the temporal requirements of any task. For instance, in perceptual decision making scenarios, the CNV reflects increasing urgency to make a decision, whereas in interval reproduction tasks, it traces anticipation of the correct time to execute the reproduction response.

In conclusion, we have demonstrated that the CNV traces urgency in both typical perceptual decision making tasks with fixed timings, as well as less commonly employed continuous monitoring tasks with temporally uncertain target onsets. We have also shown that it traces urgency in a distinct manner compared to mu/beta motor preparation, as its build-up is not influenced by sensory evidence and its termination is more closely aligned to decision commitment than response execution. Identifying these two features highlights the potential usefulness of the CNV for informing and constraining computational models, and has key implications for understanding the specific stages of the decision process that urgency acts on. Furthermore, in establishing that the CNV is an urgency signature, our results contribute to the long-standing debate surrounding its functional role.

## Methods

### Experiment 1

#### Participants

Participants were thirty young adults (14 females, mean age: 23.10 years, age range: 18-34 years). All subjects were right-handed, with normal or corrected-to-normal vision and no history of personal or familial neurological or psychiatric illness. Participants provided written consent prior to their participation. The experimental procedures were approved by the Research Ethics Committee of the School of Psychology, Trinity College Dublin in accordance with the Declaration of Helsinki. Participants were compensated with a payment of €30 for their participation. Four participants were removed due to low levels of choice accuracy (<60%), and an additional participant was removed from the analysis due to excessive artefacts (see details below), leaving 25 participants in the final sample.

#### Experimental Procedure

Participants performed the task in a dark, sound-attenuated room, seated approximately 57 cm from the computer monitor. Stimuli were presented on a dark grey background on a 51cm CRT monitor, at a 100 Hz refresh rate and screen resolution of 1024×768 pixels. All task stimuli were generated using custom made MATLAB scripts and presented via Psychtoolbox. The annulus stimulus consisted of two overlaid grating patterns (spatial frequency = 1 cycle per degree) presented in a circular aperture (inner radius = 1°, outer radius = 6°) against a dark grey background (luminance: 65.2 cd/m2). Each grating stimulus was tilted by 45° relative to the vertical midline (left tilt = -45°, right tilt = +45°). On each trial, the left and right tilted gratings were randomly assigned to flicker at either 20 Hz or 25 Hz.

Participants performed a two-alternative, forced-choice contrast discrimination task in which they had to discriminate whether the left or right tilted gratings in an annulus stimulus had a greater contrast. Participants completed 6 experimental blocks of 60 trials. Blocks were performed in alternating speed and accuracy emphasis regimes. At the start of each block, on-screen written instructions informed participants which response regime to adopt. Block order was counterbalanced across participants. At the beginning of each trial, a central fixation point was presented, followed 200ms later by the presentation of a cue instructing participants to either “Go Fast” or “Be Accurate”. This cue was presented for 500ms, after which the fixation point was displayed again for a further 500ms. Following this, the grating stimuli were presented, fading in over a period of 400ms from 0 to 50% contrast. After the end of the fade-in period, the stimulus remained at 50% contrast for a further 400ms before the target and non-target underwent antithetical changes in contrast. The target stepped up in contrast by either 10% or 16% while the nontarget stepped down in contrast by a corresponding amount. The gratings remained at the new contrast level for 1,600ms and participants were required to indicate the grating with the highest contrast by clicking the corresponding left or right mouse button with their left or right thumb.

Participants were rewarded points based on their performance. In the Accuracy Regime, correct responses were awarded 60 points irrespective of response time, while misses were awarded 0 points. Errors were punished by deducting 60 points irrespective of response time. In the Speed Regime, rewards for correct responses and punishments for incorrect responses were scaled with response time. Correct responses were awarded up to 100 points with the number of points diminishing as a function of response time at a rate of 4.8 points every 64ms (75 points per second). Consequently, correct responses that were very slow (>1,350ms) incurred minor point deductions. Errors were punished by deducting upwards of 20 points at an increasing rate of 4 points every 64ms. Consequently, on trials where participants failed to respond within the deadline (misses), 116 points were deducted. Participants received feedback on their accuracy and points scored following each trial.

### Data Analysis

#### Behavioural Analysis

Trials were sorted according to Regime (Speed vs Accuracy) and Evidence Strength (Strong vs Weak), and the effects of these variables on choice accuracy and response time (RT) was assessed using RM-ANOVAs with Regime and Difficulty as within-subject factors. Within each condition, accuracy was plotted as a function of RT (‘conditional accuracy function’, CAF) to examine how accuracy was influenced by RT.

#### EEG Acquisition and Preprocessing

Continuous EEG data were acquired using an ActiveTwo system (BioSemi) from 128 scalp electrodes, digitised at 512 Hz. Data were analysed in custom MATLAB scripts using functions from the EEGLAB ^43^ toolbox. Blinks were detected using two vertical electro-oculogram (EOG) electrodes that were placed above and below the left eye. The continuous EEG data were low-pass filtered below 35 Hz, high-pass filtered above .05 Hz and detrended. EEG data were then re-referenced offline to the average reference.

EEG data were segmented into stimulus and response-aligned epochs. Stimulus aligned epochs were extracted from 100ms before stimulus onset to stimulus offset (+2,000ms). Response-aligned epochs were measured from -800ms pre-response to 200ms post-response. All epochs were baseline-corrected relative to the interval of -500ms to -400ms from evidence onset. This baseline window was chosen, with the aid of visual inspection of the grand-average ERP waveform, to fall before the onset of the CPP but after the conclusion of evoked potentials elicited by the appearance of the stimulus during fade-in. Trials were rejected if the bipolar vertical EOG signal (upper minus lower) exceeded an absolute value of 200μV, or if any scalp channel exceeded 100μV at any time during the stimulus-aligned epoch. To avoid excessive trial loss, channels were interpolated if their individual artefact count exceeded 10% of the total number of trials. To avoid excessive channel interpolation, a maximum of 10% of the total number of channels were permitted to be interpolated for any given participant’s data. Participants were excluded from the electrophysiological analyses entirely if, following channel inspection and interpolation, more than 40% of trials were lost due to blinks and/or EEG artefacts. On this basis, one participant was excluded from the electrophysiological analyses, though their data were retained for behavioural analyses. In order to mitigate the effects of volume conduction across the scalp, single trial epochs underwent current source density (CSD) transformation ^44^, a procedure that has been used in order to minimise spatial overlap between functionally distinct EEG components ^45^.

#### EEG Signature Analysis

The CNV was measured from the average of four fronto-central electrodes that had the steepest negative slope across participants and conditions in the lead up to response (−750ms to -250ms from response). Single trial CNV data were rejected from the analyses if the amplitude prior to response exceeded +/-3 standard deviations from the within-subject mean. The CPP was measured as the average of (2-4) central-parietal electrodes centred on the maximum pre-response amplitude (−150ms to -50ms) across conditions for each participant. Single trial CPP data were rejected from the analyses if the amplitude of the CPP at response exceeded +/-3 standard deviations from the within-subject mean. The effects of speed pressure and RT on pre-evidence onset (0 to 50ms from evidence onset) and pre-response (−250ms to -150ms from response) CNV amplitude were assessed using RM ANOVAs with Regime (Accuracy vs Speed) and RT Bin (5 evenly split bins) as within-subject factors. The effect of sensory evidence strength on the build-up rate of the CNV and CPP was assessed using a RM ANOVA with Evidence Strength (Strong vs Weak) and Signal (CNV vs CPP) as within-subject factors. For this analysis, we measured build-up rate between 150ms and 350ms post-evidence onset, excluding trials with RT less than 350ms. We also multiplied the CNV slopes by -1 so that the slopes of both signals were in the same direction. Effector-selective mu/beta (10-30Hz) amplitude was measured over the contralateral and ipsilateral hemispheres by computing a Short-Time Fourier Transform (STFT) with a window size of 400ms and a step size of 50ms. The 20 Hz and 25 Hz frequencies were removed to avoid contamination from the steady-state visual evoked potential (SSVEP) emitted from the flicker rate of the stimulus. For each participant, mu/beta was measured as the average of four electrodes that exhibited the largest pre-response (−150ms to - 50ms) lateralisation index (largest difference between left and right response trials) across conditions, taken from a cluster identified in each hemisphere over the motor cortex (located at standard sites C3 and C4). Single trials were excluded from the analyses if the mu/beta lateralisation index at response exceeded +/- 3 standard deviations from the within-subject mean. The effect of speed pressure and RT on pre-evidence onset (−100ms to 100ms from evidence onset) and pre-response (−150ms to -50ms from response) mu/beta amplitude was assessed using RM ANOVAs with Regime (Accuracy vs Speed) and RT Bin (4 evenly split bins) as within-subject factors.

#### Modelling

We fit a two-choice drift diffusion model with urgency to the behavioural data. This model assumes that decision makers arrive at a judgement by gradually accumulating noisy ‘sensory evidence’ into a decision variable (‘DV’) ^5^. This accumulation process begins at a starting point (*z*), and a response is made once the accumulated evidence reaches the bound for the correct (*a*) or incorrect (*0*) response. The rate of the evidence accumulation is defined by the drift rate (*v*), which is dependent on the strength of the sensory evidence. There is also within-trial or diffusion noise (*s*), that is added to the DV at each time point. A non-decision time parameter (*Ter*) is also included to account for delays attributable to the encoding of sensory information and execution of motor responses. Urgency was added to this model via the inclusion of a 3 parameter Weibull function ^10^, that specified how the decision bounds changed as a function of time. Between-trial variability in starting point (*sz*) and drift rate (η) were also included in the model.

The model was fit separately to data from the Accuracy and Speed Regime. Following the standard approach adopted in the literature, within-trial noise was fixed to 0.1 in both Regimes. In addition, between-trial and starting point variability were fixed to 0.2 and 0.1 respectively, based on fits to the pooled data from both Regimes. All other parameters were freely estimated. Parameter values were estimated by minimising the G^2^ statistic with a particle swarm optimisation routine ^46^. The model was fit to 10 observed values, which were generated by calculating five RT quantiles (0.1, 0.3, 0.5, 0.7, 0.9) from the RT distribution on correct and incorrect trials respectively, and multiplying the proportion of trials lying between those quantiles by the total number of trials. All responses from 0ms to the response deadline were included in the fits. Expected values were calculated from the model simulations using the same method, and the goodness of fit between the expected and observed values was assessed using a G^2^ test.

### Experiment 2

#### Participants

Participants were 30 young adults (19 females, mean age: 21.50 years, age range: 19-34 years). All subjects were right-handed, with normal or corrected-to-normal vision and no history of personal or familial neurological or psychiatric illness. Participants provided written consent prior to their participation. The experimental procedures were approved by the Research Ethics Committee of the School of Psychology, Trinity College Dublin in accordance with the Declaration of Helsinki. Participants were compensated with a payment of 10 euro per hour for their participation.

#### Experimental Procedure

Participants performed the task in a dark, sound-attenuated room, seated approximately 57 cm from the computer monitor. Stimuli were presented on a black background on a 51cm CRT monitor, at a 75 Hz refresh rate and screen resolution of 1024×768 pixels. All task stimuli were generated using custom made MATLAB scripts and presented via Psychtoolbox. The random dot motion stimulus consisted of 60 white dots (each 4 pixels with a speed of 6 deg/s), presented centrally in a circular aperture with a diameter of 8 degrees, flickering at a rate of 75 Hz. A white fixation point was located at the centre of the aperture throughout stimulus presentation.

Participants performed a two-alternative, forced-choice random dot motion task in which they had to discriminate the direction (left or right) of a cloud of moving dots, reporting their choice using left and right mouse clicks with their left and right thumbs. Participants performed the task in blocks with varying levels of time pressure. In the High and Low Speed Emphasis blocks, participants were required to respond within 1200ms and 1800ms of evidence onset respectively. In Delayed Response blocks, participants were instructed to withhold reporting their choices until a response cue appeared at 1800ms. Participants performed 12 blocks of 80 trials that were separated into 4 sets of 3 (High Speed, Low Speed, Delayed Response), so that participants were not presented with more of one block type than the others in the first or second half of the experiment.

Within each set of three blocks, block order was randomised. Each trial began with a variable fixation period (400ms, 600ms, 900ms, 1200ms, 1500ms) during which only the central fixation point was presented on screen. The dot motion stimulus was then presented at 0% motion coherence for 680ms before a portion of the dots began to move to the left or right to ensure that visual potentials would have resolved prior to coherent motion onset. In the speeded response conditions, coherent motion was presented up until response deadline elapsed (1200ms for High Speed, 1800ms for Low Speed). In the Delayed Response condition, the stimulus offset at 1800ms and participants were presented with a response cue (the word ‘Respond’ appeared on screen centrally in white text), following which they had to indicate their response within 1000ms. Within each block, evidence strength randomly varied between four coherence levels (0%, 5%, 10% and 20%). At the end of each trial, participants were presented with on-screen feedback about their performance. Participants were informed if they responded prior to evidence onset (‘Too Fast’), or if they failed to respond within the response deadline (‘Too Slow’). If they responded within the correct time window, participants were informed of the accuracy of their chosen response (‘Correct’ vs ‘Incorrect’). Participants received points based on their performance: they were awarded 50 points per correct response. Feedback on the number of correct responses and points scored was delivered every 20 trials.

### Data Analysis

#### Behavioural Analysis

Trials were sorted according to Regime (High Speed, Low Speed, Delayed Response) and Evidence Strength (0%, 5%, 10%, 20%), and the effects of these variables on choice accuracy and response time (RT) was assessed using RM-ANOVAs with Regime and Evidence Strength as within-subject factors. Within each condition, accuracy was plotted as a function of RT (‘conditional accuracy function’, CAF) to examine how accuracy was influenced by RT.

#### EEG Acquisition and Preprocessing

Continuous EEG data were acquired using an ActiveTwo system (BioSemi) from 128 scalp electrodes, digitised at 512 Hz. Data were analysed in custom MATLAB using functions from the EEGLAB ^43^ toolbox. Blinks were detected using two vertical electro-oculogram (EOG) electrodes that were placed above and below the left eye. The continuous EEG data were low-pass filtered below 35 Hz, high-pass filtered above .05 Hz and detrended. Noisy channels were interpolated and the EEG data were re-referenced offline to the average reference.

EEG data were segmented into stimulus and response-aligned epochs. Stimulus aligned epochs were extracted from -750ms relative to coherent motion onset to either 1200ms, 1800ms or 2300ms post coherent motion onset for the high speed, low speed and delayed response conditions, respectively. Response-aligned epochs were measured from –700ms pre-response to 400ms post-response. All epochs were baseline-corrected relative to the interval of -750ms to -700ms prior to coherent motion onset. Trials were rejected if the bipolar vertical EOG signal (upper minus lower) exceeded an absolute value of 200μV or if any scalp channel exceeded 100μV at any time during the stimulus aligned epoch. In order to mitigate the effects of volume conduction across the scalp, single trial epochs underwent current source density (CSD) transformation ^44^, a procedure that has been used in order to minimise spatial overlap between functionally distinct EEG components ^45^.

#### EEG Signature Analysis

The CNV was measured from the average of 4 fronto-central electrodes that had the steepest negative slope across conditions for each participant in the window of -500ms to -200ms pre response, based on the free response conditions. The CPP was measured as the average of 4 central-parietal electrodes centred on the maximum pre-response amplitude (−150ms to -80ms) across conditions for each participant. As in Experiment 1, the effect of speed pressure and RT on pre-evidence onset (0ms - 50ms from evidence onset) and pre-response (−250ms to -150ms from response) CNV amplitude was assessed using RM ANOVAs with Regime (High, Low, Delayed Response) and RT Bin (5 evenly split bins) as within-subject factors. The effect of sensory evidence strength on the build-up rate of both signals was assessed using an RM ANOVA with Evidence Strength (Low vs High) and Signal (CNV vs CPP) as within-subject factors. For this analysis, we measured build-up rate between 150ms and 350ms post-evidence onset, excluding trials with RT less than 350ms. Effector-selective mu/beta (10-30Hz) amplitude was measured over the contralateral and ipsilateral hemispheres by computing a Short-Time Fourier Transform (STFT) with a window size of 400ms and a step size of 50ms. For each participant, mu/beta was measured as the average of 4 electrodes that exhibited the largest pre-response (−150ms to -50ms) lateralisation index (largest difference between left and right response trials) across conditions, taken from a cluster identified in each hemisphere over the motor cortex (located at standard sites C3 and C4). The effect of speed pressure and RT on pre-evidence onset (−100ms to 100ms from evidence onset) and pre-response (−150ms to -50ms from response) mu/beta amplitude was assessed using RM ANOVAs with Regime (High Speed, Low Speed, Delayed Response) and RT Bin (4 evenly split bins) as within-subject factors.

We examined the peak latency of the CNV in the Delayed Response to assess whether the anticipatory processes indexed by the CNV terminated at the time of choice commitment (indexed by CPP peak) or response execution. For this analysis, we exclusively used trials in the 20% coherence condition, as this was the only coherence level at which participants reliably made their decision prior to the 1800ms deadline imposed in the Low Speed condition.

### Experiment 3

#### Participants

Participants were twenty-three young adults (14 females, mean age: 22.5 years, age range: 18-29 years). All participants had normal or corrected to normal vision, had no personal or family history of epilepsy or unexplained fainting, no sensitivity to flickering light and no personal history of neurological or psychiatric illness or brain injury. Three participants were left handed. Participants provided written consent prior to their participation. The experimental procedures were approved by the Research Ethics Committee of the School of Psychology, Trinity College Dublin in accordance with the Declaration of Helsinki. Participants were compensated with a €20 gift voucher for their participation. Participants were also informed that, once data had been collected from all participants, they would each be given a bonus voucher of up to €10 based on their performance. However, all participants received a €10 voucher.

#### Experimental Procedure

Participants performed the task in a dark, sound-attenuated room, seated approximately 75 cm from the computer monitor. Stimuli were presented on a black background on a 51cm CRT monitor, at a 85 Hz refresh rate and screen resolution of 1024×768 pixels. All task stimuli were generated using custom made MATLAB scripts and presented via Psychtoolbox. The random dot motion stimulus consisted of 118 white dots (each 4×4 pixels with a speed of 3.33 deg/s), presented centrally in a circular aperture with a diameter of 8 degrees, flickering at a rate of 17 Hz. A white, square fixation (5×5 pixels) was located in the centre of the aperture at all times.

Participants performed a continuous monitoring version of the random dot motion task ^45^ in which they monitored a cloud of randomly moving dots for intermittent ‘targets’, defined by a step change from incoherent to coherent leftward or rightward motion for 2 seconds. During incoherent motion, dots were displayed in a new, random location within the circular aperture on each frame. Coherent motion was generated by displacing a portion of the dots by a distance of 0.24 degrees in either the left or right direction on the next frame. Targets were displayed at two coherence levels (14% and 20%). There were three inter-target-intervals (ITIs), during which the dots moved incoherently (3.6s, 6.6s, 8.4s). The coherence level and ITI for each trial varied pseudo-randomly within each block. Participants held the mouse in both hands and used left and right thumb presses to indicate the direction of motion.

Participants performed the task in alternating speed and accuracy regime blocks. The regime was manipulated by task instructions and points structure. In Accuracy blocks, participants were instructed to take their time and respond as accurately as possible. In Speed blocks, participants were instructed to prioritise responding as fast as possible. Participants were rewarded points based on their performance. In accuracy blocks, they were awarded 40 points if they made a correct response during target presentation (2 seconds). In Speed blocks, participants had just under half as much time to earn points for correct responses (900ms). This deadline was selected based on pilot testing that indicated that median response times were circa 1 second without speed emphasis. To compensate for the increased difficulty of this shortened deadline and ensure participants did not become demotivated, participants were rewarded 70 points for correct responses in speed blocks. If participants responded correctly after the deadline but during target presentation (i.e. between 900 and 2000ms, labelled as ‘Too Late’ responses rather than misses), they received 0 points. In both conditions, participants were penalised for missing targets (−30 points) and making false alarms (−20 points). Participants received feedback on the points scored, number of misses, and number of false alarms at the end of each block. On speed blocks, participants also received feedback on the number of Too Late responses they had made). Each participant performed 8 blocks (4 accuracy, 4 speed), with 48 targets presented in each block.

Prior to the experiment, participants performed a series of short practice blocks (12 targets). In the first block, participants were presented with coherent motion set at 80%, and verbalised the direction of the targets. Participants then completed one practice block at 20% coherence and one practice block at 14% coherence. In these blocks, participants were instructed to respond as quickly and accurately as possible. Some participants completed additional practice blocks at 14% coherence, until the experimenter was satisfied that they could detect targets at this coherence level. Finally, participants were informed of the speed-accuracy manipulation, and performed a short block of the speed condition to become familiar with the additional feedback they received for Too Late responses at the end of speed blocks.

### Data Analysis

#### Behavioural Analysis

Trials were sorted according to Regime (Accuracy, Speed), Coherence (Low (14%), High (20%)), and ITI (Short (3.6s), Medium, (6.6s), Long (8.4s)). Four behavioural output measures were analysed: response time (RT), miss rate, false alarm rate and choice accuracy. RT was calculated as the time from coherent motion onset to response. Miss rate was calculated as the proportion of total targets that were missed. False alarm rate was calculated as the proportion of total trials in which the participant responded during incoherent motion. The effects of ITI on RT, Miss rate, and False Alarm were analysed using repeated measures ANOVA with ITI as a within-subjects factor.

#### EEG Acquisition and Preprocessing

Continuous EEG data were acquired using an ActiveTwo system (BioSemi) from 128 scalp electrodes, digitised at 512 Hz. Data were analysed in custom MATLAB and Python 3 scripts, using functions from the EEGLAB ^43^ and MNE-python ^47^ toolboxes. Blinks were detected using two vertical electro-oculogram (EOG) electrodes that were placed above and below the left eye. EEG data were low-pass filtered offline at 30Hz. No offline high-pass filter was applied. EEG data for each block were detrended to remove slow linear drifts. Noisy channels were interpolated and the EEG data were re-referenced offline to the average reference.

To examine EEG activity over the ITI period, epochs were extracted from 200ms after target offset until the onset of the next target. These epochs were baseline-corrected to the period from 200ms to 400ms from ITI onset and any trials that had a response in this baseline period were removed. To perform artefact rejection on the long ITI epochs without excessive trial loss, these epochs were separated into 8 one second chunks (starting at 400ms and ending at 8400ms). For artefact rejection alone, each chunk was re-baselined to the -200ms to 0ms period prior to its onset. On a given chunk, if more than 12 scalp channels (approximately 10% of channels) exceeded an absolute value of 100μV, or EOG activity exceeded 200μV, that chunk was rejected. If there were no EOG artefacts, and the number of scalp electrodes with an artefact for all chunks on a given trial was less than 12, these electrodes were interpolated for all chunks on the trial. Following artefact rejection, EEG data were transformed into Current Source Density (CSD) using the CSD toolbox in MATLAB ^44^. One participant was removed from the analysis due to excessive artefacts, leaving 23 participants for the analysis (13 females, mean age: 22.47 years, age range: 18-29 years).

#### EEG Signature Analysis

The CNV was measured from two fronto-central electrodes centred on the minimum grand-average pre-response amplitude (located between standard sites Fz and FCz). To assess motor preparation over the course of the ITI, we measured effector-selective mu/beta (10-30Hz) amplitude by computing a Short-Time Fourier Transform (STFT) with a window size of 300ms (3 full cycles of the lowest mu/beta frequency) and a step size of 25ms. The 17Hz frequency was omitted to avoid contamination from the steady-state visual evoked potential (SSVEP) emitted from the flicker rate of the stimulus. Mu/beta amplitude was measured from three electrodes over the motor cortex on each hemisphere, centred on the standard sites C3 and C4. To compare the time courses of the CNV and motor preparation over the ITI, mu/beta amplitude was entered into a linear regression predicting CNV amplitude. For this analysis, the CNV was first downsampled to the same number of timepoints as mu/beta. We performed these analyses on both the grand average across Regimes (reported main text) and separately within Regime (reported in S1).

## Acknowledgments

This work was funded by an SFI research grant to R.G.O. and S.P.K. (19/US/3599). R.G.O. was funded also by the European Research Council Consolidator Grant IndDecision – 865474, and S.P.K. also by a Wellcome Trust Investigator Award (219572/Z/19/Z).

## Author Contributions

J.D, E.McN and H.McC collected the data. C.A.D & H.McC preprocessed and analysed the behavioural and EEG data, H.McC & R.G.O performed the statistical analyses, D.P.McG performed the computational modelling. H.McC and R.G.O drafted the article. All authors contributed substantially to discussion of the analyses conducted and/or edited the manuscript before submission.

