## Supplementary Materials for "A Movement-Independent Signature of Urgency During Human Perceptual Decision Making"

### S1: CNV over the ITI as a function of Regime

Regression analysis reported in Experiment 3 performed separately on each Regime indicated that the CNV and Mu/Beta in both Regimes followed a highly similar time course over the ITI (Accuracy: $R^2 = 0.81$, $F(1, 302) = 1296$, $p < 0.001$, $r = 0.90$; Speed: $R^2 = 0.81$, $F(1, 302) = 1298$, $p < 0.001$, $r = 0.90$).

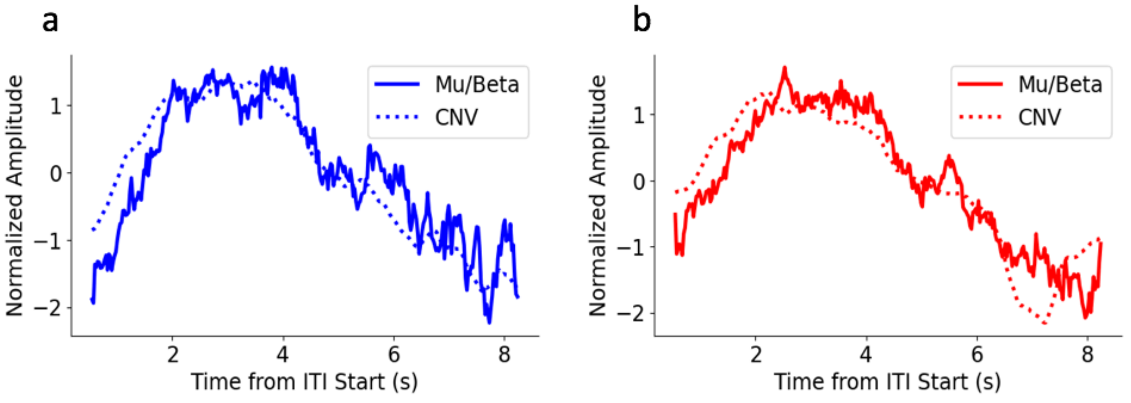


*Figure S1. Mu/Beta and CNV during the ITI in the Accuracy (a) and Speed (b) Regime.*
